# Investigation of biological activities of *Colocasia gigantea* Hook.f. leaves and PASS prediction, in silico molecular docking with ADME/T analysis of its isolated bioactive compounds

**DOI:** 10.1101/2020.05.18.101113

**Authors:** Safaet Alam, Nazim Uddin Emon, Mohammad A. Rashid, Mohammad Arman, Mohammad Rashedul Haque

## Abstract

**Background:** *Colocasia gigantea* is locally named as kochu and also better known due to its various healing power. This research is to investigate the antidiarrheal, antimicrobial, and antioxidant possibilities of the methanol soluble extract of *Colocasia gigantea*.

**Methods:** Antidiarrheal investigation was performed by using *in vivo* castor oil induced diarrheal method where as *in vitro* antimicrobial and antioxidant investigation have been implemented by disc diffusion and DPPH scavenging method respectively. Moreover, *in silico* studies were followed by molecular docking analysis of several secondary metabolites were appraised with Schrödinger-Maestro v 11.1.

**Results:** The induction of plant extract (200 and 400 mg/kg, b.w, p.o), the castor oil mediated diarrhea has been minimized 19.05 % (p < 0.05) and 42.86 % (p < 0.001) respectively. The methanolic extract of *C. gigantea* showed mild sensitivity against almost all the tested strains but it shows high consistency of phenolic content and furthermore yielded 67.68 μg/mL of IC_50_ value in the DPPH test. The higher and lower binding affinity was shown in beta-amyrin and monoglyceryl stearic acid against the kappa-opioid receptor (PDB ID: 4DJH) with a docking score of -3.28 kcal/mol and -6.64 kcal/mol respectively. In the antimicrobial investigation, Penduletin and Beta-Amyrin showed the highest and lowest binding affinity against the selected receptors with the docking score of -8.27 kcal/mol and -1.66 kcal/mol respectively.

**Conclusion:** The results of our scientific research reflect that the methanol soluble extract of *C. gigantea* is safe which may provide possibilities of alleviation of diarrhea and as a potential wellspring of antioxidants which can be considered as an alternate source for exploration of new medicinal products.

## Introduction

Diarrhea, a common disease in the tropical countries and can be interpreted as an incidence of daily stool exceeding 200 gm comprised of 60 to 95% of water. Infants and children suffer from diarrhea most, and mortality from diarrhea is high compared with other diseases. (Sini, Umar et al. 2008). Unhygienic living style plays a key role in making people lying to diarrhea. The World Health Organization (WHO) declared diarrhea as the second leading cause for child mortality under five years of age (Lutterodt 1992) which is almost 1.87 million reflecting 19 % of total child deaths (Boschi-Pinto, Velebit et al. 2008). In cases of an immune-suppressed condition like human immunodeficiency virus/acquired immune deficiency syndrome (HIV/AIDS), patients become more vulnerable with concurrent chronic diarrhea (Ahmed, McGaw et al. 2014). Diarrhea which is a common gastrointestinal disorder with latent fatal implications can impose serious public health hazards (Page, Hustache et al. 2011). Study protocol of diarrhea in human and animal models reveals several triggering etiologies including infections, food intolerances, intestinal disorders, and medications (Zhang, Wang et al. 2013). Various enteropathogens like *Escherichia coli, Shigella flexneri, Salmonella typhi Staphylococcus aureus*, and *Candida albicans* are major causative agents that can provoke diarrhea (Konaté, Yomalan et al. 2015). Despite establishing a diarrhea disease control program (DDC) by the World Health Organization (WHO), diarrhea is still alarming in developing countries (Wansi, Nguelefack-Mbuyo et al. 2014). This program includes the practice of traditional medicines along with evaluation of health education and prevention approaches predominantly based on herbal constituents (Konaté, Yomalan et al. 2015).

As pathogenic bacterias result in a major cause of morbidity and mortality in humans, Pharmaceutical companies are determined to produce masses of new antibacterials which are becoming significant against infections and drawing global concern (Djeussi, Noumedem et al. 2013). A couple of justifications make clinical microbiologists interested in antimicrobials from plant extracts including the possibility of phytochemicals to be the arsenal of antimicrobial agents prescribed by the physicians and making people aware of the risks with the misuse of traditional antibiotics (Sher 2009). Additionally, antibiotic resistance pops up the window of considering intensive research to preserve the effectiveness of currently available antibiotic agents along with findings newer classes (Guo-Hong, Pei-Ji et al. 2004) as plant-derived natural products exhibit improved shield against microbial attacks (Dixon 2001).

Oxidative stress, induced by oxygen radicals, is believed to be a primary factor in various deteriorating diseases, such as cancer (Muramatsu, Kogawa et al. 1995) atherosclerosis (Steinberg, Parthasarathy et al. 1989), gastric ulcer (Das, Bandyopadhyay et al. 1997), and other conditions. There is a rising interest in natural antioxidants which are derived from plants as bioactive components. The protective acts of diets rich in fruits and vegetables against cardiovascular diseases and certain cancers have been attributed partly due to the antioxidant contains including therein, particularly to phenolic compounds (Ferreira, Barros et al. 2007). The roles of free radicals and active oxygen in the treatment of human diseases against cancer, aging, and atherosclerosis have been documented (Aruoma, Murcia et al. 1993). Electron acceptors, such as molecular oxygen, react rapidly with free radicals to become radicals themselves, and also referred to as reactive oxygen species (ROS). The ROS includes superoxide anions (O_2_), hydrogen peroxide (H_2_O_2_), and hydroxyl radicals (OH) (Horton 2003). The importance of the antioxidant constituents of plant materials in the maintenance of health and protection from coronary heart disease and cancer is also raising attention among the scientists, food manufacturers, and consumers as the trend of the future is moving toward functional food with specific health effects (Kumaran and Karunakaran 2007). Many antioxidant compounds, naturally occurring from plant sources, have been identified as a free radical or active oxygen scavengers (Zheng and Wang 2001). Recently, interest has increased considerably in finding naturally occurring antioxidants to use in foods or medicinal materials to replace synthetic antioxidants, which are being restricted due to their side effects such as carcinogenicity (Kumaran and Karunakaran 2007).

Healing with medicinal plants is as old as mankind itself (Petrovska 2012). Natural products derived from plants for the treatment of diseases have proved that nature stands as a golden mark to show the interrelationship between man and the environment (Oladeji 2016). Since time immemorial the history of humankind has availed plant extracts of various plants to treat and heal their physical agony (Mukul, Uddin et al. 2007) and from that ancient time plant and its extracts play a pivotal role and confers therapeutic importance in world health care up to today (Yadav, Kumar et al. 2006). 80% of drug substances are either a direct derivative of the natural component or a refined version of the natural part of the plant extracts (Maridass and De Britto 2008).

The genus *Colocasia* is represented by 13 species worldwide (Yin 2006) among which eight species were found in Asia and the Malay Archipelago initially (Mayo, Bogner et al. 1997). In Bangladesh, so far this genus contains the following nine species: *C. gigantea* (Blume) Hook. f., *C. fallax* Schott, *C. affinis* Schott, *C. esculenta* (L.) Schott, *C. oresbia* A. Hay, *C. heterochroma* H. Li et Z.X. & Wei, *C. virosa* Kunth, *C. lihengiae* C.L. Long et K.M. Liu, and *C. mannii* Hook. f.(Ara 2007). *Colocasia* is a flowering plant genus under Araceae family native to southeastern Asia and the Indian subcontinent which are widely cultivated and naturalized in other tropical and subtropical regions (Wagner, Herbst et al. 1990). Genus of *Colocasia* leaves has demonstrated the potentiality of antidiabetic, antihypertensive, immunoprotective, neuroprotective, and anticarcinogenic activities (Gupta, Kumar et al. 2019). *Colocasia gigantea* (Family: Araceae) is a perennial herb of 1.5-3 m tall available in South-East Asia and leaf stalk is consumed as a vegetable (Ivancic, Roupsard et al. 2008). *C. gigantea* is abundantly found in Bangladesh and locally known as Kochu. This plant is also known as giant elephant ear or indian taro. Phytochemical extraction and structure elucidation of *Colocasia* leaves yield chemical compounds such as isoorientin, orientin, isoschaftoside, Lut-6-C-Hex-8-C-Pent, vicenin, alpha-amyrin, beta-amyrin, mono glycerol stearic acid, penduletin anthraquinones, apigenin, catechins, cinnamic acid derivatives, vitexin, and isovitexin (Liu, Liu et al. 2014, Gupta, Kumar et al. 2019).

With technological advancements and better opportunities to explore the previously inaccessible sources of natural products will provide many promising drug candidates to unleash unexplored opportunities. Therefore, to reveal new chapters of phytochemicals pharmaceutical industries are trying to find new natural lead compounds to perform pre-clinical and eventually clinical trials to ensure efficacy and safety as it is believed that natural moieties are blessed with fewer side effects. However, considering all auspicious factors, this study is conducted to evaluate the antidiarrheal, antimicrobial, and antioxidant activity of methanolic extract of *C. gigantea* by biological and computational approaches.

## Material and method

### Collection and extraction of plant

The leaves of *C. gigantea* were collected from Bandarban in May 2019. The plant was identified at the Bangladesh National Herbarium (BNH) and a voucher specimen (Accession no: BNH-58077) has been deposited for this collection. After proper washing, the whole pant was sun-dried for several days. The dried plant was then grounded to a coarse powder by using a high capacity grinding machine. 800 gm of the powdered material was taken in a clean, and round bottom flask (5 liters) and soaked in 2.4 liters of methanol. The container with its content was kept for a period of 10 days accompanying daily shaking and stirring. The whole mixture was then filtered through a fresh cotton plug and finally with a Whatman No.1 filter paper. The volume of the filtrate was then reduced through using a Buchi Rotavapor at low temperature and pressure. The weight of the crude extract was found 60.82 gm.

### Drugs and chemicals

All drugs and chemicals used in this research were of analytical grade. Methanol, Tween-80, was purchased from Merck (Darmstadt, Germany). 1,1-diphenyl-2-picrylhydrazyl radical (DPPH), Folin-Ciocalteau reagent (FCR) were obtained from Sigma Chemicals Co. (St. Louis, MO, USA). Loperamide (Square Pharmaceuticals Ltd., Dhaka, Bangladesh); ciprofloxacin (Beximco Banglades Ltd, Dhaka, Bangladesh); were procured from the mentioned sources.

### Experimental animals

Swiss-albino mice of either sex, aged 4-5 weeks, acquired from the Animal Resource Branch of the International Centre for Diarrhoeal Diseases and Research, Bangladesh (ICDDR,B) and were used for the experiment. They were housed in standard polypropylene cages and kept under controlled room temperature (24 ± 2 °C; relative humidity 60-70 %) in a 12 h light-dark cycle and fed ICDDR,B formulated rodent food and water (*ad libitum*). As these animals are very sensitive to environmental changes, they were kept in the environment where the experiment will take place before the test for at least 3-4 days. These studies were held following the internationally accepted principle for appropriate use of laboratory animals, namely the National Institutes of Health (NIH) and International Council for Laboratory Animal Science (ICLAS).

### In vivo acute toxicity test

The oral toxicity test was conducted under normal conditions in laboratories and followed the “Organization for Environmental Control Development” guidelines (OECD: Guidelines 420;) fixed-Dose Method (Lipnick, Cotruvo et al. 1995) and after the administration of high oral doses to the mice, several parameters were recorded throughout the 72 hours. As a result, there was no lethality, no behavioral change (sedation, excitability), or no allergies reaction was appeared after the oral administration methanol soluble leaves extract of *C. gigantea*.

### *In vivo* castor oil induced diarrhea

The anti-diarrheal activity of the methanolic extract of leaves of *C. gigantea* was evaluated using the method of castor oil-induced diarrhea in mice (Ezeja, Ezeigbo et al. 2012). According to this method, each mouse was fed with 1 mL of the highly pure analytical grade of castor oil which would induce diarrhea. The numbers of fecal stools were recorded for each mouse. The observations of the experimental groups were compared against that of the control to evaluate the anti-diarrheal activity of the samples. The animals were divided into control, positive control, and test groups containing six mice in each group. Control group received vehicle (1% Tween 80 in water) at dose 10 mL/kg orally. The positive control group received loperamide at the dose of 2 mg/kg orally. The test group received a methanolic extract of leaves of *C. gigantea* the doses of 200 and 400 (mg/kg, b.w; p.o). Each animal was placed in an individual cage; the floor lining was changed at every hour. Diarrhea was induced by oral administration of castor oil to each mouse after the above treatment. During an observation period of 4 hours; the number of diarrhoeic feces excreted by the animals was recorded.

### *In vitro* antibacterial assay

The antimicrobial assessment has been performed by following the disc diffusion method (Karaman, Şahin et al. 2003). In this classical method, on nutrient agar medium uniformly seeded with the test microorganisms dried and sterilized filter paper discs (6 mm diameter) containing the test samples of known amounts are placed. Antibiotics diffuse from a confined source through the nutrient agar gel and create a concentration gradient. Standard antibiotic (Amoxicillin) discs and blank discs are used as positive and negative control. To allow maximum diffusion of the test materials to surround media these plates are kept at low temperature (4 °C) for 16 to 24 hours (Barry, 1976). For optimum growth of the organisms, the plates are then inverted and incubated at 37°C for 24 hours. The test materials having antimicrobial property inhibit microbial growth in the media surrounding the discs and thereby yield a clear, distinct area defined as a zone of inhibition. The diameter of the zone of inhibition expressed in millimeter is then measured to determine the antimicrobial activity of the test agent (Mahboubi and Haghi 2008). Antimicrobial activity was evaluated on the clinically isolated strains of following pathogens *Bacillus cereus, Bacillus megaterium, Bacillus subtilis, Staphylococcus aureus, Sarcina lutea* as gram-positive bacteria *and Escherichia coli, Pseudomonas aeruginosa, Salmonella paratyphi, Salmonella typhi, Shigella dysenteriae, Vibrio mimicus*, and *Vibrio parahemolyticus* as a gram-negative bacteria. Wells of 6 mm diameter were punched into the agar medium with sterile cork borer under aseptic conditions and filled with 50 μL of 250 mg/mL of plant extract, solvent blank, and standard antibiotic. Serial dilutions were prepared from 250 mg/mL of the plant extract using DMSO to make 250, 125, 62.5, 31.25, and 15.625 mg/mL. The wells were inoculated with 0.1 mL aliquot of test organisms (106 CFU/mL) having serial dilutions of the extract (50 μL, each). Standard Amoxicillin (30 μg/disc) discs were used as a positive control to ensure the activity of standard antibiotics against the test organisms as well as for comparison of the response produced by the known antimicrobial agent with that of produced by the test sample.

### *In vitro* total phenolic contents analysis

The total phenolic content of *C. gigantea* extractives was measured employing the method as described (Blainski, Lopes et al. 2013) involving Folin-Ciocalteu reagent as an oxidizing agent and gallic acid as standard. To 0.5 mL of extract solution (2 mg/mL), 2.5 mL of Folin-Ciocalteu reagent (diluted 10 times with water) and 2.0 mL of Na_2_CO_3_ (7.5 % w/v) solution was added. The mixture was incubated for 20 minutes at room temperature. After 20 minutes the absorbance was measured at 760 nm by UV-spectrophotometer and using the standard curve prepared from gallic acid solution with different concentrations, the total phenols content of the sample was measured. The phenolic contents of the sample were expressed as mg of GAE (gallic acid equivalent)/gm of the extract.

### *In vitro* assay of free radical scavenging activity

DPPH was used to evaluate the free radical scavenging activity (antioxidant potential) of various compounds and medicinal plants (Choi, Jhun et al. 2000). 2.0 mL of a methanol solution of the sample (extractives/control) at different concentration (500 μg/mL to 0.977 μg/mL) were mixed with 3.0 mL of a DPPH methanol solution (20 μg/mL). After a 30 min reaction period at room temperature in a dark place, the absorbance was measured at 517 nm against methanol as blank by UV spectrophotometer. Inhibition of free radical DPPH in percent (%) was calculated as follows

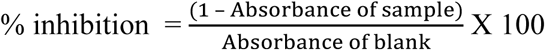

Where A_blank_ is the absorbance of the control reaction (containing all reagents except the test material).

### Molecular Docking Analysis: Selection of compounds for the computational study

Alpha-amyrin, beta-amyrin, monoglyceryl stearic acid and penduletin were selected based on the availability as major compounds through chemical investigation. The chemical structure of isolated compounds has been retrieved from PubChem (https://pubchem.ncbi.nlm.nih.gov/) and presented in **Figure 1**.

**Figure 1.**
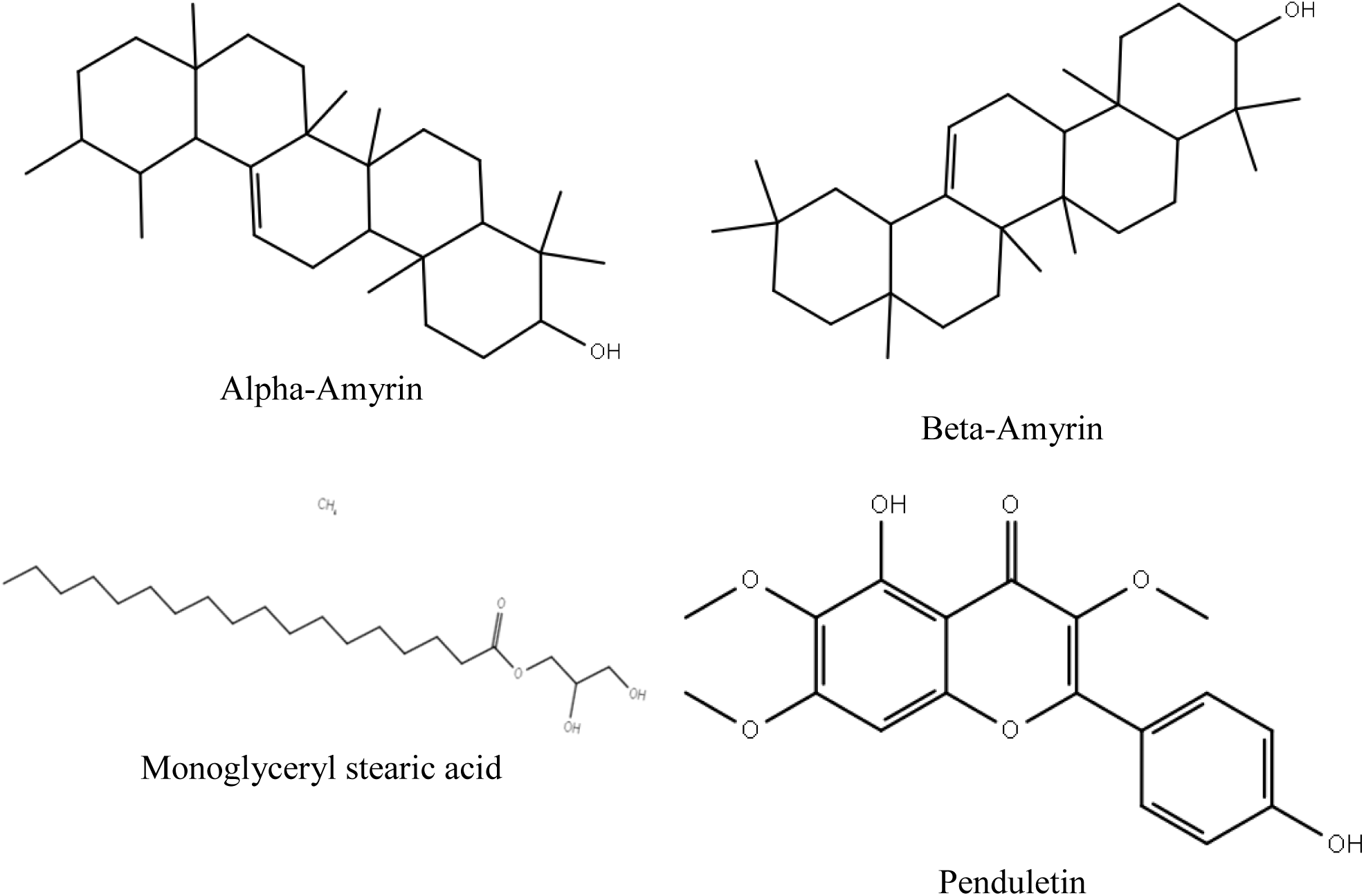
Chemical structures of Alpha-Amyrin, Beta-Amyrin, Monoglyceryl stearic acid and Penduletin used for the computational study.

### Molecular Docking Analysis: Ligand Preparation

The chemical structures of the four compounds (alpha-amyrin, beta-amyrin, monoglyceryl stearic acid, and penduletin) of *C.gigantea* were downloaded from the PubChem compound database (https://pubchem.ncbi.nlm.nih.gov/). By means of the Lig Preptool which was incorporated in Schrödingersuite-Maestro v 11.1, the ligand was created where the following factors were used as follows: neutralized at pH 7.0 ± 2.0 using Epik 2.2 and the OPLS_2003 force field were used for minimization.

### Molecular Docking Analysis: Enzyme/Receptor Preparation

In the present study, an orally active drug should fulfill the following drug-likeness parameters to demonstrate their pharmaceutical fidelity such as moles have been obtained from the Protein Data Bank RCSBPDB (Prlić, Bliven et al. 2010). kappa-opioid receptor (PDB: 4DJH and 6VI4) (Wu, Wacker et al. 2012, Che, English et al. 2020), human delta-opioid receptor (PDB: 4RWD) (Fenalti, Zatsepin et al. 2015), human glutamate carboxypeptidase II (PDB: 4P4D) (Novakova, Cerny et al. 2016), Beta-ketoaryl-ACP synthase 3 receptor (PDB: 1HNJ) (Qiu, Janson et al. 2001) Reca mini intein-zeise’s salt (PDB: 5K08) (Chan 2020), and Aromatic Prenyl Transferase (PDB: 1ZB6) (Kuzuyama, Noel et al. 2005). The enzyme/receptor was prepared for a docking experiment using Protein Preparation Wizard, which embedded in Schrödinger suite-Maestro v 11.1.

### Molecular Docking Analysis: Glide Standard Precision Docking

A molecular docking study was performed to reveal the possible mechanism of action of the selected compounds behind the biological activities of the *C. gigantea* against the respective enzymes/receptor for an antidiarrheal, and antibacterial activity. Docking experiments were performed using Glide standard precision docking, which was embedded in Schrödingersuite-Maestro v 11.1, as we described previously (Emon, Jahan et al. 2020).

### In Silico Study: Determination of Pharmacokinetic Parameters by SwissADME

The pharmacokinetic parameters or drug-likeness properties of the selected compounds were determined by SwissADME online total molecular weight of the compounds, Lipophilicity (LogP), the number of hydrogen-bond acceptors, and the number of hydrogen-bond donors based on the Lipinski’s rule.

### In Silico Study: Toxicological Properties Prediction by Admet SAR

The toxicological properties of the designated compounds have been determined by the admetSAR online tool (http://lmmd.ecust.edu.cn/admetsar1/predict/) since toxicity is a prime apprehension throughout the development of new drugs. The present study projected Ames toxicity, carcinogenic properties, and acute rat toxicity.

### PASS prediction study

The four major phytoconstituent Alpha-Amyrin, Beta-Amyrin, Monoglyceryl stearic acid Penduletin were investigated for evaluating the antidiarrheal, antibacterial, and antioxidant activities by using PASS online program.

### Statistical Analysis

The data was presented as a standard error mean (SEM). Statistical analyses using single-way ANOVA were conducted and followed by Dunnett’s multiple comparison tests. The observed values were compared to the control group and were considered statistically significant at p < 0.05, p < 0.01, p < 0.001.

## Results

### Castor oil induced diarrheal assay

The methanolic extract of *C. gigantea* exhibited promising anti-diarrheal activity with a 19.05 % (p < 0.05) and 42.86 % (p < 0.001) reduction of diarrhea at the dose of 200 mg/kg and 400 mg/kg compared to the standard loperamide 68.18% which has extremely statistically significant anti-diarrheal activity. The result of this study shows that the methanolic extract of *C. gigantea* possesses noteworthy anti-diarrheal activity which is dose dependent and activity is statistically significant (p < 0.001) at 400 mg/kg body weight dose. The findings have been shown in **Figure 2**.

**Figure 2.**
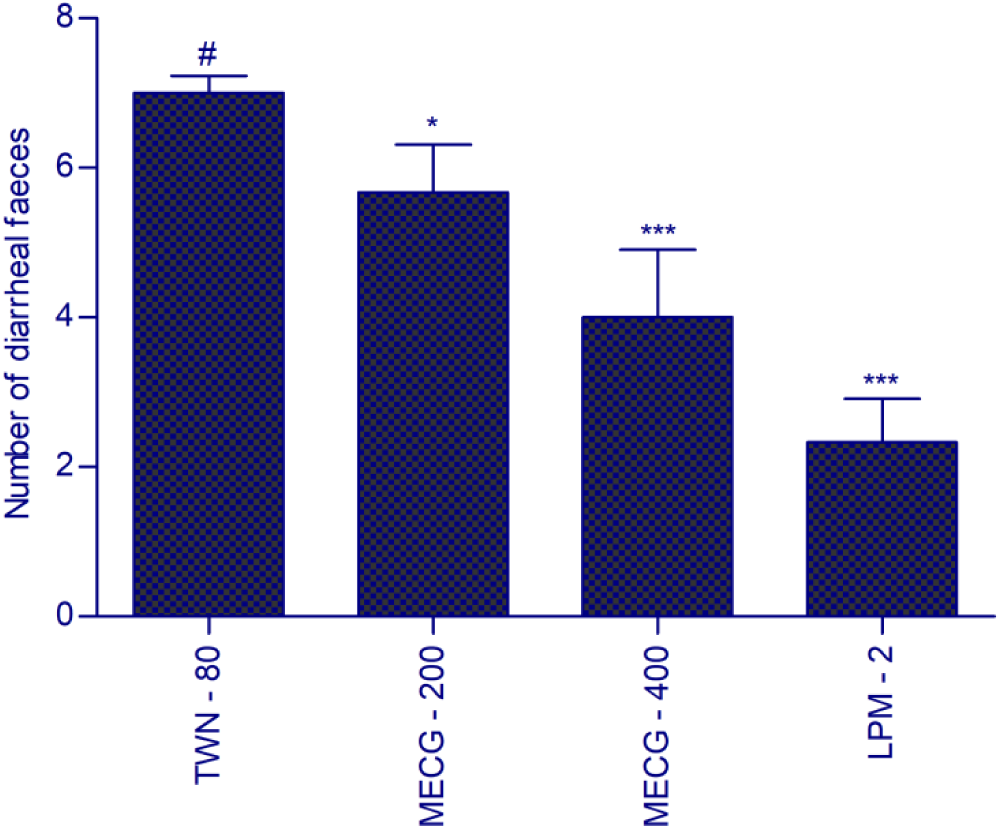
Effect of methanolic extract of leaves of *C. gigantea* on castor oil induced diarrhea in mice. Each value has been expressed as mean ± SEM (n = 6). \**p* < 0.05, **p < 0.01, ***p < 0.001 and compared with the control group (Dunnett’s test). TWN-80 = 1% tween – 80, MECG = methanol extract of *C. gigantea* leaves, LPM-2 = Loperamide 2 mg/kg.

### Analysis of Antimicrobial Activity

The Methanol Soluble Extract exhibit mild inhibition against microbial growth having a zone of inhibition ranged from 11 mm to 18 mm. the maximum zone of inhibition produced by *C.gigantea* was found to be 18 mm against *Staphylococcus aureus* and *Salmonella typhi* followed by 17 mm against *Vibrio parahemolyticus* respectively. Besides, Amoxicillin used as standard drug exhibited prompt antimicrobial activity from 41 mm against *Staphylococcus aureus* and *Pseudomonas aerginosa*. The growth of inhibition of the microbe has been presented in **Table 1.**

**Table 1.**
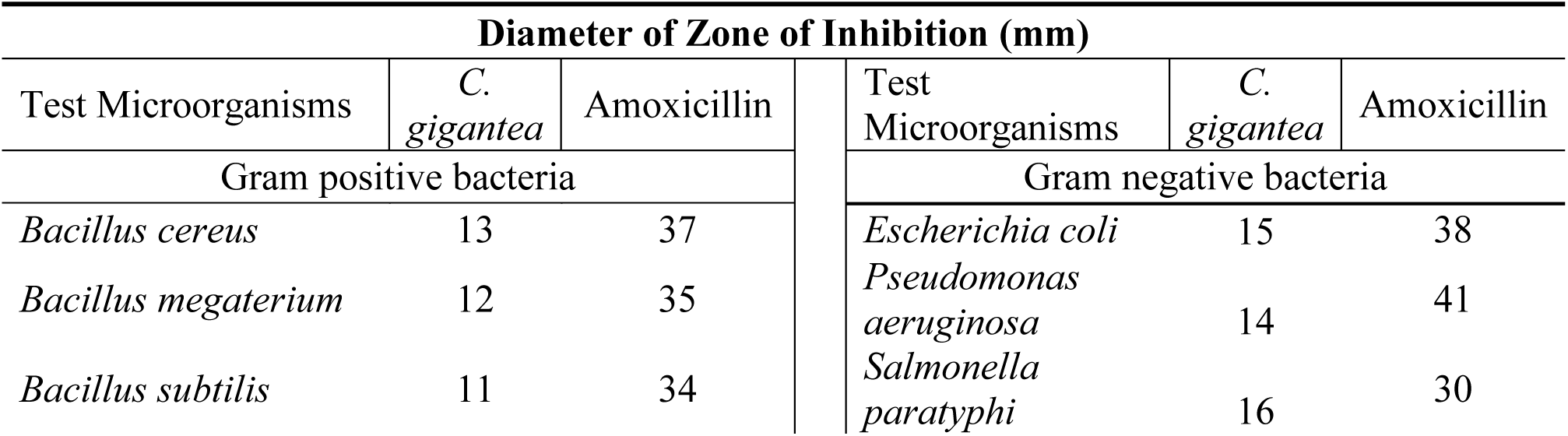
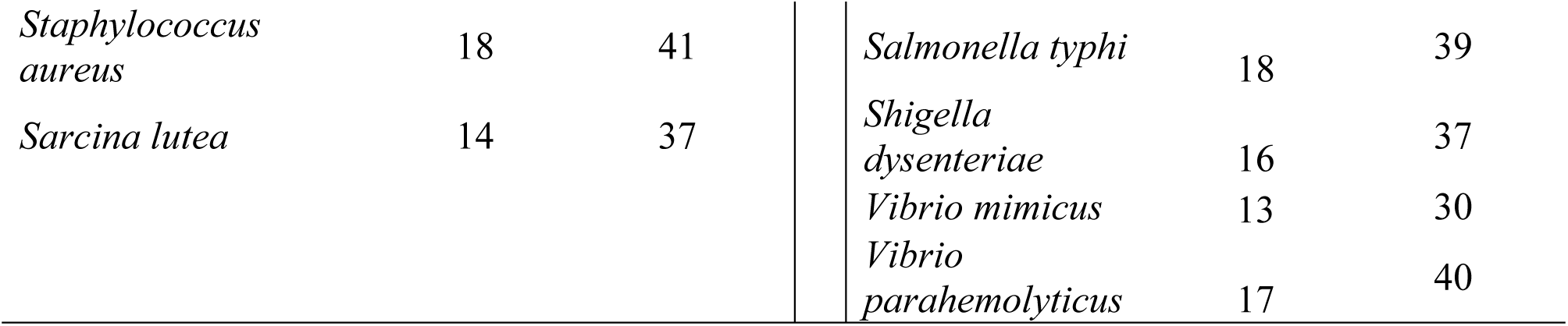
Antimicrobial activity of test samples of *C. gigantea* and Amoxicillin against gram positive and gram negative bacterial strains.

| Diameter of Zone of Inhibition (mm) |  |  |  |  |  |
| --- | --- | --- | --- | --- | --- |
| Test Microorganisms | *C. gigantea* | Amoxicillin | Test Microorganisms | *C. gigantea* | Amoxicillin |
| Gram positive bacteria |  |  | Gram negative bacteria |  |  |
| *Bacillus cereus* | 13 | 37 | *Escherichia coli* | 15 | 38 |
| *Bacillus megaterium* | 12 | 35 | *Pseudomonas aeruginosa* | 14 | 41 |
| *Bacillus subtilis* | 11 | 34 | *Salmonella paratyphi* | 16 | 30 |

|  |  |  |  |  |  |  |
| --- | --- | --- | --- | --- | --- | --- |
| *Staphylococcus aureus* | 18 | 41 |  | *Salmonella typhi* | 18 | 39 |
| *Sarcina lutea* | 14 | 37 |  | *Shigella dysenteriae* | 16 | 37 |
|  |  |  |  | *Vibrio mimicus* | 13 | 30 |
|  |  |  |  | *Vibrio parahemolyticus* | 17 | 40 |

### Total phenolic content determination of *C. gigantea*

The amount of total phenolic content for the methanolic extract of *C. gigantea* leaves has been found 38.76 mg of GAE/gm of extractives. The phenolic contents of the sample were expressed as mg of GAE (gallic acid equivalent)/gm of the extract. The average phenolic content of standard (Gallic acid) and methanolic extract of *C. gigantea* has been shown in **Table 2** and **Figure 3**.

**Table 2.** Total phenolic content determination of methanolic extract of *C. gigantean.*

| Sample | Absorbance 1 | Absorbance 2 | Average Absorbance | (mg/g of extract) |
| --- | --- | --- | --- | --- |
| *C. gigantea* | 1.59 | 1.52 | 1.55 | 38.76 |

**Figure 3.**
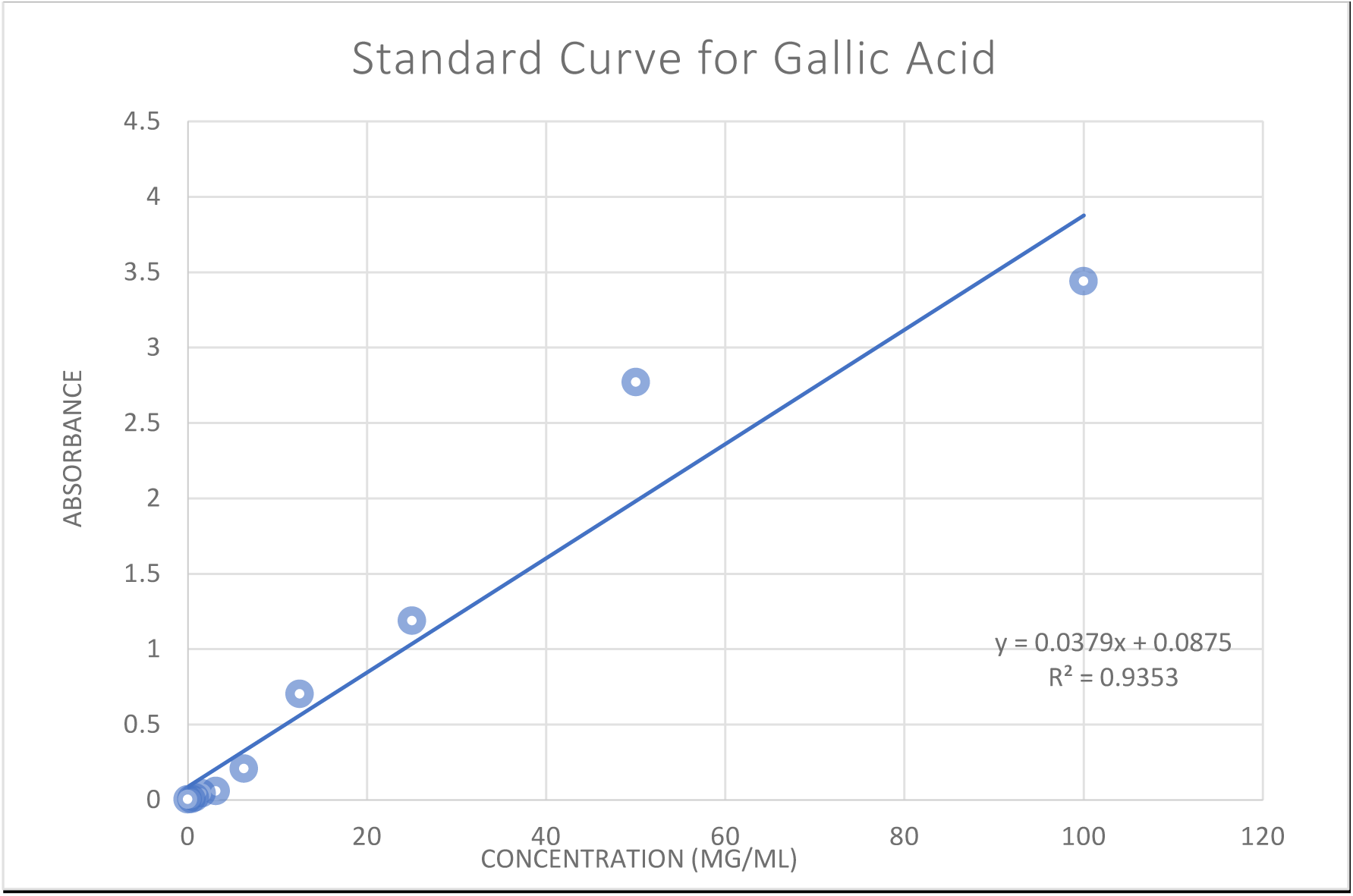
The curve of gallic acid for total phenolic contents determination of extract.

### Free radical scavenging activity (DPPH)

The IC_50_ values of methanol extract of *C. gigantea* in the DPPH method have been compared to tert-butyl-1-hydroxytoluene (BHT) and the IC_50_ value has been found 67.68 μg/mL for the methanol soluble extract of *C. gigantea.* IC_50_ value for BHT was found 20.07 μg/mL. The summary of the IC_50_ values of the test samples and the tert-butyl-1-hydroxytoluene (BHT) has been presented in **Figures 4** and **5**.

**Figure 4.**
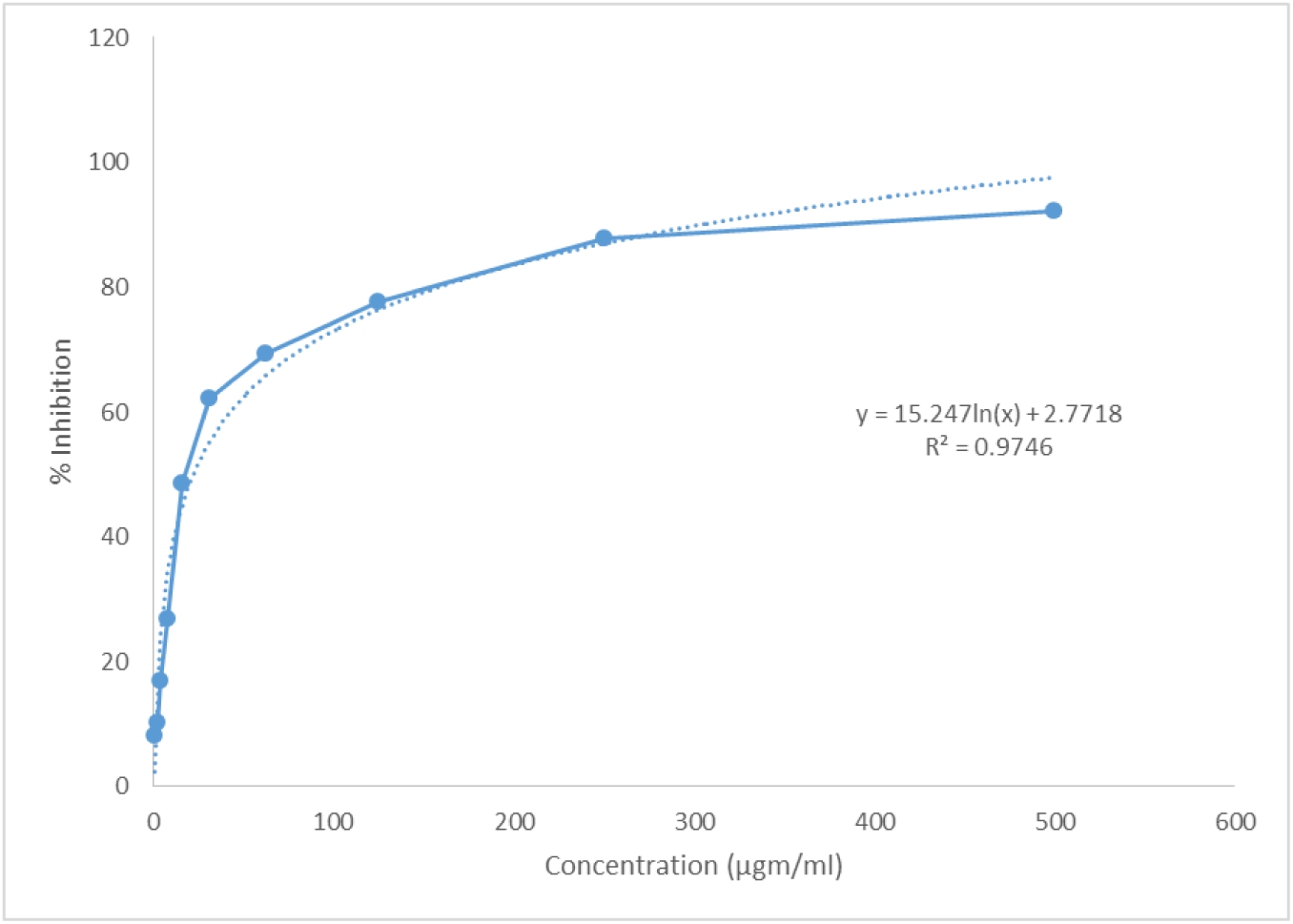
IC50 value of tert-butyl-1-hydroxytoluene (BHT).

**Figure 5.**
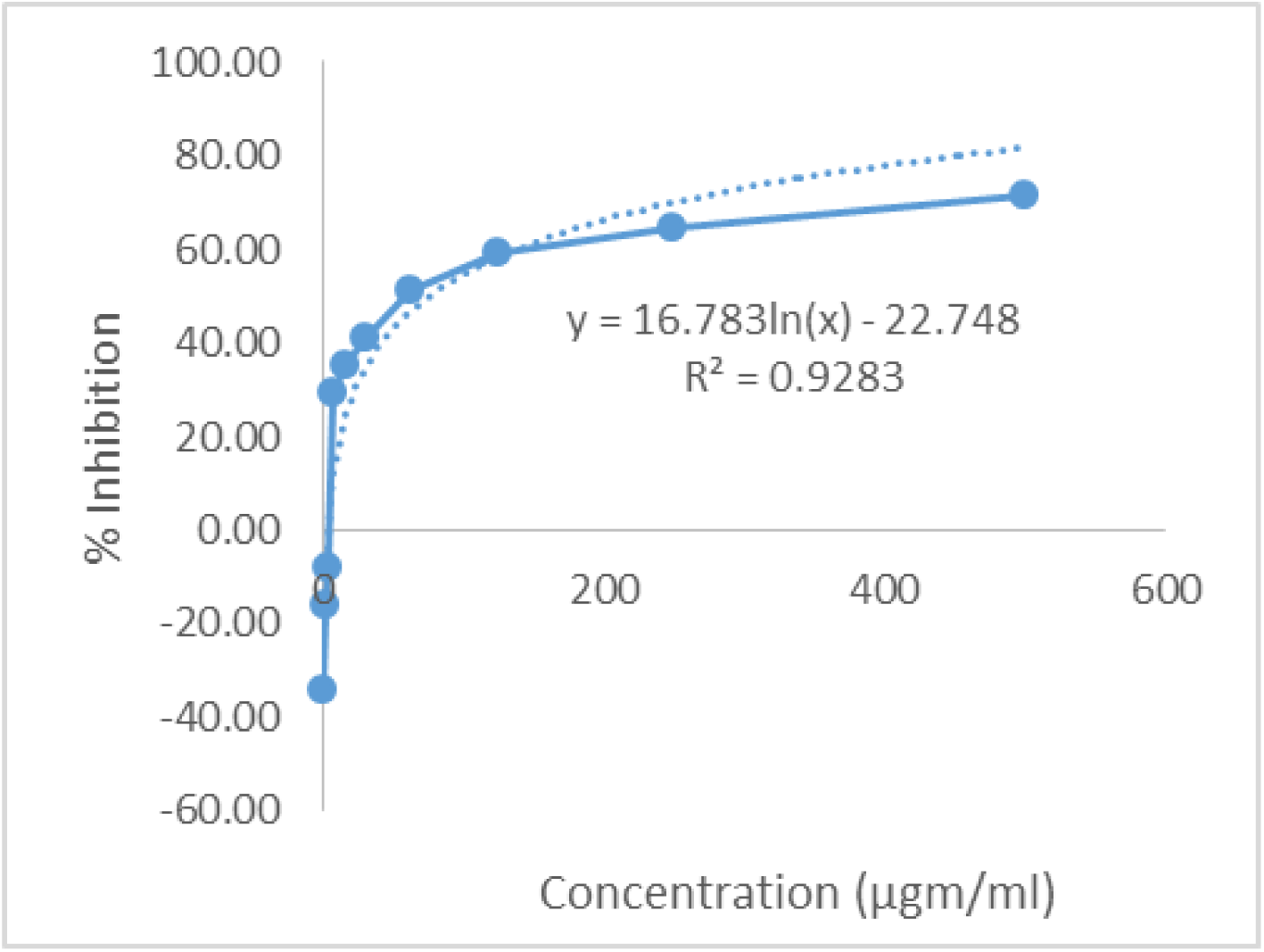
IC50 value of methanol soluble extract of *C. gigantea* observed with DPPH.

### Molecular docking study for antidiarrheal and antibacterial activity

In this case, beta-amyrin and monoglyceryl stearic acid have demonstrated the maximum and lowermost binding affinity against the kappa-opioid receptor (PDB ID: 4DJH) with a docking score of -3.28 kcal/mol and -6.64 kcal/mol respectively. The ranking order of the docking score is presented below: Monoglyceryl stearic acid > Penduletin > Alpha-Amyrin > Beta-Amyrin. Human kappa-opioid receptor (PDB ID: 6VI4) varies with a docking score of -4.03 to -6.24 (kcal/mol) respectively. The ranking order of the docking score is presented as follows: Monoglyceryl stearic acid > Alpha-Amyrin > Penduletin > Beta-Amyrin. Human delta-opioid receptor (PDB ID: 4RWD) fluctuates with a docking score of -3.88 to -5.51 (kcal/mol) respectively. The ranking order of the docking score is presented as follows: Alpha-Amyrin > Penduletin > Monoglyceryl stearic acid > Beta-Amyrin. Loperamide (reference drug) exposed - 6.64, -5.14, and -6.24 (kcal/mol) binding affinity kappa and delta opioid receptor respectively (PDB ID: 4DJH, 6VI4,and 4RWD). On the other hand, in the antimicrobial investigation, Penduletin and Beta-Amyrin displayed the highest and lowest binding affinity against the selected receptors with the docking score of -8.27 kcal/mol and -1.66 kcal/mol, respectively. human glutamate carboxypeptidase II (PDB: 4P4D) fluctuates with a docking score of -5.05 to - 6.76 (kcal/mol) respectively. The ranking order of the docking score is presented as follows: Penduletin > Alpha-Amyrin > Beta-Amyrin > Monoglyceryl stearic acid. The ranking order of the docking score of Beta-ketoaryl-ACP synthase 3 receptor (PDB: 1HNJ) with the selected compounds are presented as follows: Penduletin > Monoglyceryl stearic acid > Beta-Amyrin > Alpha-Amyrin. The ranking order of the docking score of Reca mini intein-zeise’s salt (PDB: 5K08) with the selected compounds are presented as follows: Penduletin > Alpha-Amyrin > Monoglyceryl stearic acid > Beta-Amyrin. The ranking order of the docking score of Aromatic Prenyl Transferase (PDB: 1ZB6) with designed bioactive components has been presented as follows: Penduletin > Monoglyceryl stearic acid > Alpha-Amyrin > Beta-Amyrin. Amoxicillin (reference drug) revealed -7.66, -5.23, -4.26, -6.06 (kcal/mol) binding affinity with the human glutamate carboxypeptidase II (PDB: 4P4D), Beta-ketoaryl-ACP synthase 3 receptor (PDB: 1HNJ), Reca mini intein-zeise’s salt (PDB: 5K08), and Aromatic Prenyl Transferase (PDB: 1ZB6) respectively. The results of the docking study are shown in **Table 3** and **Figure 6**.

**Table 3.**
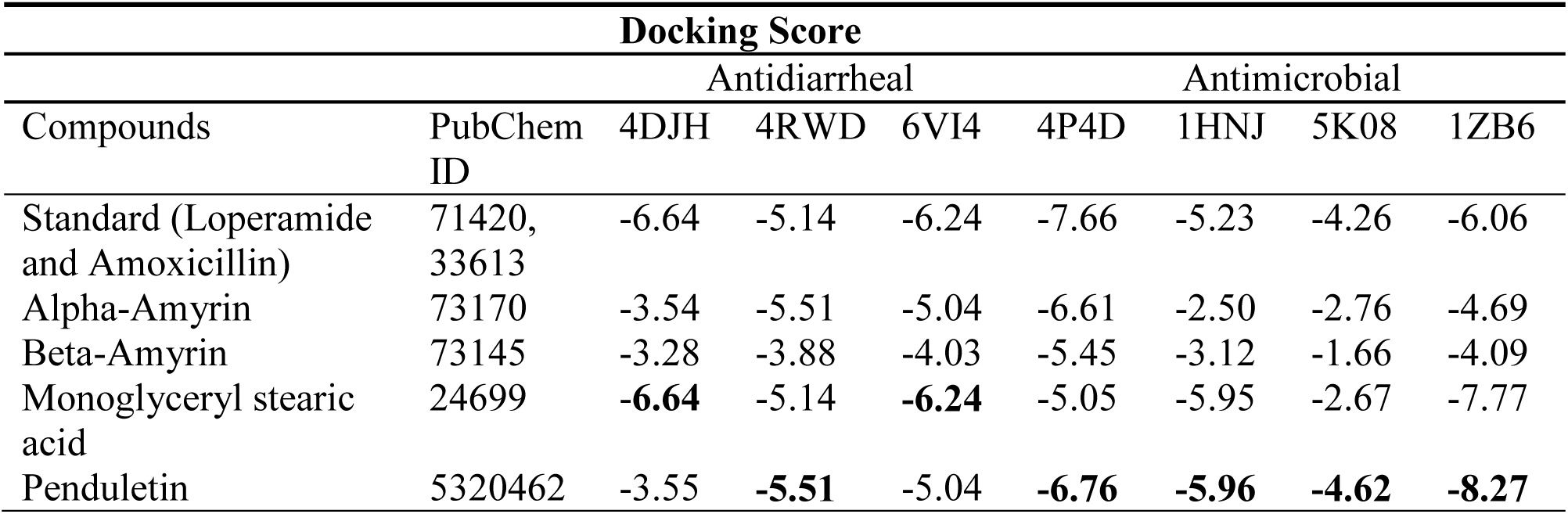
Docking scores and binding interactions of the selected compounds with the kappa-opioid receptor (PDB: 4DJH and 6VI4), human delta-opioid receptor (PDB: 4RWD), human glutamate carboxypeptidase II (PDB: 4P4D), Beta-ketoaryl-ACP synthase 3 receptor (PDB: 1HNJ), Reca mini intein-zeise’s salt (PDB: 5K08), and Aromatic Prenyl Transferase (PDB: 1ZB6) for antidiarrheal and antibacterial activity respectively.

**Figure 6.**
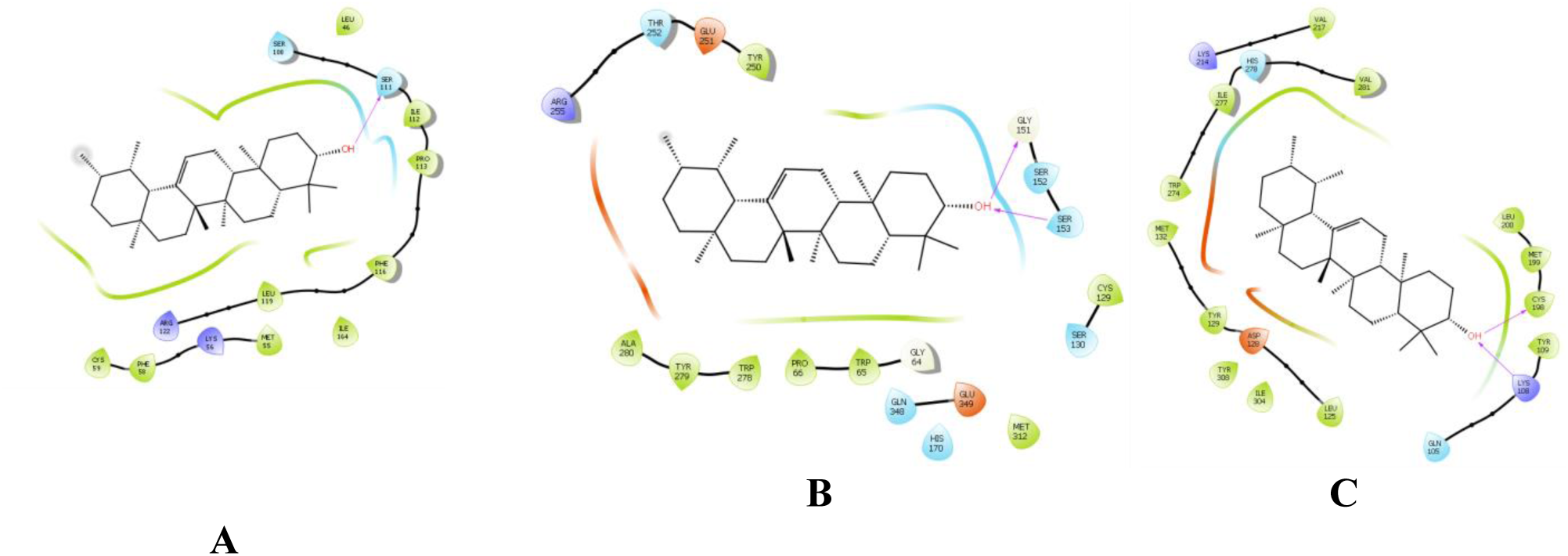
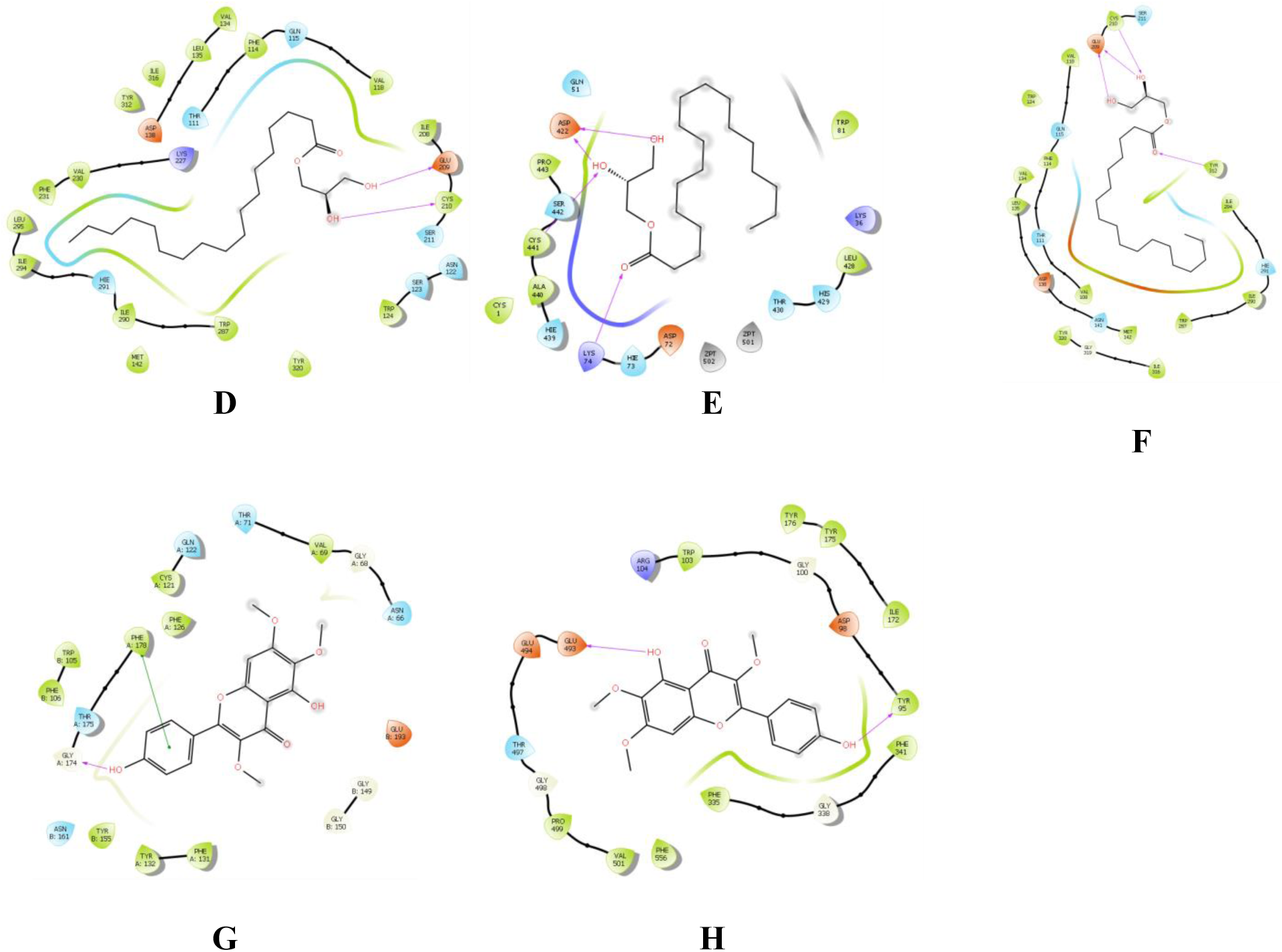
2D representation of the interactions between the best pose found for (A, B, C) Alpha-Amyrin with PDB: 4P4D, 4MR7, 4RWD, (D, E, F) Monoglyceryl stearic acid with 4DJH, 6VI4, PDB: 5K08 and (G, H) Penduletin with 4RWD and 1ZB6.

### Pharmacokinetic (ADME) and toxicological properties

Prediction The pharmacokinetic features of the substances chosen by Lipinski were determined using SwissADME, the online tool. Lipinski has here declared that if a drug/compound follows the following criteria such as molecular weight < 500 amu, Hydrogen bond donor sites < 5, Hydrogen bond acceptor sites < 10, and Lipophilicity value LogP ≤ 5, then the compound would be orally bioavailable. The study showed that all the compounds complied with the rules of Lipinski, suggesting the strong oral bioavailability of these compounds **(Table 4)**. Besides, the admetSAR online server predicted the toxicological properties of the four selected compounds. The analysis revealed that the selected compounds are non-Ames toxic, non-carcinogenic, and had low rat toxicity values.

**Table 4.** Physicochemical and toxicological properties of the compounds for good oral bioavailability.

| Compound<br>s | Molecular<br>Weight<br>(M.W)<br>(g/mol) | H-bond<br>Don<br>or | H-bond<br>Accepto<br>rs | Lipophilic<br>ity - log P<br>(o/w) | GI<br>Absorpti<br>on | AMES<br>Toxici<br>ty | Carcinoge<br>ns | Rat<br>Acute<br>Toxici<br>ty |
| --- | --- | --- | --- | --- | --- | --- | --- | --- |
| Loperamide | 477.04 | 1 | 3 | 4.13 | High | No | No | 3.6560 |
| Amoxicillin | 365.40 | 4 | 6 | 1.22 | Low | No | No | 1.7036 |
| Alpha-Amyrin | 426.72 | 1 | 1 | 4.77 | Low | No | No | 2.0842 |
| Beta-Amyrin | 426.72 | 1 | 1 | 4.75 | Low | No | No | 2.0842 |
| Monoglyceryl stearic<br>acid | 358.56 | 2 | 4 | 4.59 | High | No | No | 0.8172 |
| Penduletin | 344.32 | 2 | 7 | 2.84 | High | No | No | 3.1579 |

### PASS prediction study

Four major selected compounds of *C. gigantea* were studied by the PASS online tool for antidiarrheal, antibacterial and antioxidant activities and the potent displayed higher Pa value than Pi **(Table 5)**.

**Table 5.** PASSS prediction of standard drug and selected bioactive compounds of *C.gigantea.*

| Biological<br>Activity | Chemical Constituents |  |  |  |  |  |  |  |  |  |
| --- | --- | --- | --- | --- | --- | --- | --- | --- | --- | --- |
|  | Standard<br>(Loperamide<br>Amoxicillin<br>and Ascorbic<br>acid) |  | Alpha-Amyrin |  | Beta-Amyrin |  | Monoglyceryl<br>stearic acid |  | Penduletin |  |
|  | Pa | Pi | Pa | Pi | Pa | Pi | Pa | Pi | Pa | Pi |
| Antidiarrheal | 0.574 | 0.003 | 0.085 | 0.011 | 0.353 | 0.023 | 0.754 | 0.003 | 0.588 | 0.006 |
| Antibacterial | 0.761 | 0.003 | 0.200 | 0.117 | 0.206 | 0.112 | 0.291 | 0.06 | 0.359 | 0.041 |
| Antioxidant | 0.928 | 0.003 | 0.411 | 0.011 | 0.405 | 0.01 | 0.276 | 0.028 | 0.714 | 0.004 |

## Discussion

Several steps including detection and characterization of bioactive substances are required to address the therapeutic activity of medicinal plants (Coan, Ottl et al. 2011). Even without a clear concept of proper dosing profile, indigenous people exploit medicinal plants in various forms including pastes, juices, or boiled leaf extracts which make researchers curious about exposing the pharmacologic potentiality by the efficient solvent extraction process. Bioactive compounds involve several compounds like flavonoids, alkaloids, tannins, alkaloids, terpenoids, and many other classes that are promptly miscible in methanol due to high polarity of 5.1 along with few non-polar constituents too. This justifies widely use of methanol in the extraction process of medicinal plants which is also used in the investigation of *C. gigantea* too (Boeing, Barizão et al. 2014). Several studies on the treatment of a number of pathological disorders including diarrhea, microbial, and oxidative degradations have been conducted using noble molecules with lower side effects. This proximate study includes the pharmacological assessment of methanolic extract of *C. gigantea* maneuvering isolated bioactive phytochemicals to aim antidiarrheal, antimicrobial, and antioxidant effects followed by computational analysis (In silico molecular docking, ADME and toxicity predictions) of it’s isolated bioactive compounds.

In this study, *C. gigantea* was found significant (p < 0.05 and 0.001) dose-dependent inhibition of the frequency of diarrheal feces. Lack of harmony between intestinal smooth muscle motility and/or absorption pattern of the GI tract can inaugurate diarrhea (Gidudu, Sack et al. 2011). The usage of Castor oil as a diarrhea medication has been well recognized (Shiferie and Shibeshi 2013). Castor oil, a very nifty laxative used as diarrhea inducer which is hydrolyzed in the upper small intestine to ricinoleic acid and able to provoke fluid secretion, inhibit water and electrolyte absorption, reduce active Na+ and K+ absorption, and decrease Na+, K+, - ATPase in the small intestine and colon (Ammon, Thomas et al. 1974) is accomplished by the irritant effect ricinoleic acid liberated by pancreatic acid (Mbagwu and Adeyemi 2008). Prostaglandins possessing a basic role in the pathophysiology of diarrhea can be released by ricinoleic acid that regulates the gastrointestinal tract, stimulate motility secretion and eventually causes diarrhea (Agunu, Yusuf et al. 2005, Hu, Gao et al. 2009). Recently revealed molecular mechanism suggests activation of EP3 prostanoid receptor by ricinoleic acid which mediates pharmacologic effects of castor oil. Intestinal and uterine-muscle cells have been triggered by ricinoleic acid via EP3 prostanoid receptors which illuminate the cellular and molecular mechanism of inducing laxative outcome of castor oil. The antidiarrheal property of methanolic extract of *C. gigantea* may be exhibited via several mechanisms including the reduction of prostaglandin secretions (Tunaru, Althoff et al. 2012). Plant extract containing flavonoids and alkaloids modifies cyclo-oxygenase 1 and 2 (COX-1, COX-2) and lipo-oxygenase (LOX) production which obstruct prostaglandin and autacoids production (Mishra, Seth et al. 2016). Flavonoids, a large group of polyphenolic compounds holding a wide variety of biological effects such as antioxidant, anti-inflammatory, antispasmodic, and antidiarrheal activities (Qin, Wang et al. 2011) can display antidiarrheal activity too by restricting intestinal motility and hydro-electrolytic secretions (Meite, N’guessan et al. 2009).

Antimicrobial property by an extract depends on the phytochemical composition, extracting solvent, effective solubility and miscibility of the active component in the test medium, the vulnerability of the test organisms, and the method used in evaluation (Ahmed, McGaw et al. 2014). Previous research suggests a bunch of phytochemical compounds like glycoside, saponin, tannin, flavonoids, terpenoid, and alkaloids as antimicrobial entities (Mendonça-Filho 2006). This study was propagated to assess the antimicrobial effect of *C. gigantea* and from the investigation, it was noticed that *C. gigantea* is mildly susceptible to the test strains. Therefore, it is an unambiguous inspection that *C. gigantea* can be considered as a source of antimicrobial moieties for further researches.

Plant extracts rich in polyphenols which can exhibit redox properties by absorbing and neutralizing free radicals utilizing scavenging properties of their hydroxyl group can demonstrate antioxidant activity (Hatano, YAsUHARA et al. 1990, Kähkönen, Hopia et al. 1999). Among those, the flavonoid is considered as supremely competent scavengers of most oxidizing molecules, including quenching single and triplet oxygen or decomposing peroxides and various free radicals implicated in several disease conditions (Saeed, Khan et al. 2012). *C. gigantea* showed a concentration-dependent antiradical activity by inhibiting DPPH radical with an IC_50_ value of 67.68 μg/mL and the total phenolic compound was determined 38.76 mg of GAE/ gm of crude extract. The result promotes that *C. gigantea* possesses dose-dependent hydrogen donating capabilities and acts as a prominent source antioxidant. By the end of enumerating all upshots, our denouement suggested that alkaloids, phenol, flavonoids, terpenoids, tannins, etc; maybe the major contributors to the antidiarrheal, antibacterial, and antioxidant activities of methanol soluble leaves extract of *C. gigantea*.

Molecular docking analyses were employed extensively in the estimation of ligand-target relationships and to gain a deeper understanding of the biological activity of natural products. It provides more insights into probable mechanisms of action and binding mode within the binding pockets of several proteins (Khan, Nazir et al. 2019). Four isolated compounds within *C.gigantea* have been selected for docking tests to provide greater insight into the biological activity (antidiarrheal and antibacterial). These compounds were then docked against seven targeted receptors, namely the kappa opioid receptor (PDB: 4DJH and 6VI4), human delta-opioid receptor (PDB: 4RWD), human glutamate carboxypeptidase II (PDB: 4P4D), Beta-ketoaryl-ACP synthase 3 receptor (PDB: 1HNJ), Reca mini intein-zeise’s salt (PDB: 5K08), and Aromatic Prenyl Transferase (PDB: 1ZB6). Docking score revealed that, among the four compounds, seven interacted with several amino acid residues through hydrogen and docking scores ranging between -1.66 to -8.27 kcal/mol. Specifically, selected bioactive compounds of *C.gigantea* directly interacted with a number of amino acid residues within the kappa opioid receptor (PDB: 4DJH and 6VI4), and human delta-opioid receptor (PDB: 4RWD) with docking scores ranging from -3.28 to -6.64 kcal/mol. From these results, we can conclude that the studied phytoconstituents may in part be responsible for the antidiarrheal activity of *C.gigantea* through interaction with these target proteins. In addition, in the antimicrobial investigation, the four compounds were docked with, human glutamate carboxypeptidase II (PDB: 4P4D), Beta-ketoaryl-ACP synthase 3 receptor (PDB: 1HNJ), Reca mini intein-zeise’s salt (PDB: 5K08), and Aromatic Prenyl Transferase (PDB: 1ZB6) and displayed docking scores varying from -1.66 to - 8.27 kcal/mol. From the results, it is clear that the phytoconstituent Penduletin displayed the highest score against Aromatic Prenyl Transferase, followed by Monoglyceryl stearic acid, Beta-Amyrin, Alpha-Amyrin.

The online prediction software ADME was also used to investigate the drug-like properties, pharmacokinetics, and physicochemical characteristics in all bioactive compounds. According to Lipinski’s law, all the bioactive compounds exhibited orally active drug likeness properties. Compounds with lower molecular weight, lipophilicity, and hydrogen bonding are stated to be highly permeable (Duffy, Devocelle et al. 2015), good absorption, and bioavailability (Lipinski, Lombardo et al. 1997).

To assess the possible pharmacological profile of the compounds we used the structures based prevision system for biological activity, namely Prediction for Activity Spectra for Substances (PASS). The consequences recommended numerous activities, among these, we have found possible activity values (Pa range 0.085 – 0.754) for all four compounds for antidiarrheal, antibacterial, and antioxidant actions, supporting our laboratory investigations of methanolic extract of *C.gigantea*. In addition, the wider ability for the species has been predicted by many other experiments. In summary, we support and encourage the use of *C.gigantea* through our detailed analyzes using complementary methods. The effects observed may be caused by the combined behavior of many phytoconstituents, both those reported herein and other compounds that have not yet been described.

## Funding

This research received no funding and hereby the study was conducted by the self-fundings of the author.

## Acknowledgment

The research was supported by the Department of Pharmaceutical Chemistry, Faculty of Pharmacy, University of Dhaka, Dhaka-1000, Bangladesh and Department of Pharmacy, International Islamic University Chittagong, Chittagong-4318, Bangladesh. The authors of the manuscript want to show their gratitude to the authority of the respective department.

## Conflict of Interest

Authors of the manuscript declare no conflict of interest.

## Ethical consideration

The current study procedures were reviewed and approved by the “P&D committee’ of the Department of Pharmaceutical Chemistry, Faculty of Pharmacy, University of Dhaka, Bangladesh, with reference number: Pharm-P&D-7/19-163.

## Authors Contribution

SA conceptualized and designed the study protocol. Along with SA, NUE prepared the plant extract, experimented the designed protocols, collected the data, calculated the data. NUE, SA, and MA designed and conducted the computational analysis. NU and SA drafted the manuscript. DMAR and DMRH supervised, monitored the research, and revised the final draft manuscript. All authors read and approved the final manuscript for publication.

## Notes

### Competing Interest Statement

The authors have declared no competing interest.

## Reference

Agunu, A., S. Yusuf, G. O. Andrew, A. U. Zezi and E. M. Abdurahman (2005). “Evaluation of five medicinal plants used in diarrhoea treatment in Nigeria.” Journal of Ethnopharmacology 101(1-3): 27–30.

Ahmed, A. S., L. J. McGaw, N. Moodley, V. Naidoo and J. N. Eloff (2014). “Cytotoxic, antimicrobial, antioxidant, antilipoxygenase activities and phenolic composition of Ozoroa and Searsia species (Anacardiaceae) used in South African traditional medicine for treating diarrhoea.” South African journal of botany 95: 9–18.

Ammon, H., P. Thomas and S. Phillips (1974). “Effects of oleic and ricinoleic acids on net jejunal water and electrolyte movement. Perfusion studies in man.” The Journal of clinical investigation 53(2): 374–379.

Ara, H. (2007). “Araceae In: Siddiqui.” KU, Islam, MA, Ahmed, ZU, Begum, ZNT, Hassan, MA, Khondker, M., Rahman, MM, Kabir, SMH, Ahmad, M., Ahmed, ATA, Rahman, AKA and Haque, EU (eds.). Encyclopedia of Flora and Fauna of Bangladesh 11: 19–98.

Aruoma, O. I., A. Murcia, J. Butler and B. Halliwell (1993). “Evaluation of the antioxidant and prooxidant actions of gallic acid and its derivatives.” Journal of Agricultural and Food Chemistry 41(11): 1880–1885.

Blainski, A., G. C. Lopes and J. C. P. De Mello (2013). “Application and analysis of the Folin Ciocalteu method for the determination of the total phenolic content from Limonium Brasiliense L.” Molecules 18(6): 6852–6865.

Boeing, J. S., É. O. Barizão, B. C. e Silva, P. F. Montanher, V. de Cinque Almeida and J. V. Visentainer (2014). “Evaluation of solvent effect on the extraction of phenolic compounds and antioxidant capacities from the berries: application of principal component analysis.” Chemistry Central Journal 8(1): 48.

Boschi-Pinto, C., L. Velebit and K. Shibuya (2008). “Estimating child mortality due to diarrhoea in developing countries.” Bulletin of the World Health Organization 86: 710–717.

Chan, H., Pearson, S., Green, C.M., Li, Z., Zhang, J., Lippard, S., Belfort, G., Shekhtman, A., Li, H.M., Belfort, M. (2020). “RecA mini intein-Zeise’s salt complex.”

Che, T., J. English, B. E. Krumm, K. Kim, E. Pardon, R. H. J. Olsen, S. Wang, S. Zhang, J. F. Diberto, N. Sciaky, F. I. Carroll, J. Steyaert, D. Wacker and B. L. Roth (2020). “Nanobody-enabled monitoring of kappa opioid receptor states.” Nature communications 11(1): 1145–1145.

Choi, H. Y., E. J. Jhun, B. O. Lim, I. M. Chung, S. H. Kyung and D. K. Park (2000). “Application of flow injection—chemiluminescence to the study of radical scavenging activity in plants.” Phytotherapy Research: An International Journal Devoted to Pharmacological and Toxicological Evaluation of Natural Product Derivatives 14(4): 250–253.

Coan, K. E., J. Ottl and M. Klumpp (2011). “Non-stoichiometric inhibition in biochemical high-throughput screening.” Expert opinion on drug discovery 6(4): 405–417.

Das, D., D. Bandyopadhyay, M. Bhattacharjee and R. K. Banerjee (1997). “Hydroxyl radical is the major causative factor in stress-induced gastric ulceration.” Free Radical Biology and Medicine 23(1): 8–18.

Dixon, R. A. (2001). “Natural products and plant disease resistance.” Nature 411(6839): 843–847.

Djeussi, D. E., J. A. Noumedem, J. A. Seukep, A. G. Fankam, I. K. Voukeng, S. B. Tankeo, A. H. Nkuete and V. Kuete (2013). “Antibacterial activities of selected edible plants extracts against multidrug-resistant Gram-negative bacteria.” BMC complementary and alternative medicine 13(1): 164.

Duffy, F. J., M. Devocelle and D. C. Shields (2015). Computational approaches to developing short cyclic peptide modulators of protein–protein interactions. Computational Peptidology, Springer: 241–271.

Emon, N. U., I. Jahan and M. A. Sayeed (2020). “Investigation of antinociceptive, anti-inflammatory and thrombolytic activity of Caesalpinia digyna (Rottl.) leaves by experimental and computational approaches.” Advances in Traditional Medicine: 1–9.

Ezeja, I. M., I. I. Ezeigbo, K. G. Madubuike, N. E. Udeh, I. A. Ukweni, S. C. Akomas and D. C. Ifenkwe (2012). “Antidiarrheal activity of Pterocarpus erinaceus methanol leaf extract in experimentally–induced diarrhea.” Asian Pacific Journal of Tropical Medicine 5(2): 147–150.

Fenalti, G., N. A. Zatsepin, C. Betti, P. Giguere, G. W. Han, A. Ishchenko, W. Liu, K. Guillemyn, H. Zhang, D. James, D. Wang, U. Weierstall, J. C. H. Spence, S. Boutet, M. Messerschmidt, G. J. Williams, C. Gati, O. M. Yefanov, T. A. White, D. Oberthuer, M. Metz, C. H. Yoon, A. Barty, H. N. Chapman, S. Basu, J. Coe, C. E. Conrad, R. Fromme, P. Fromme, D. Tourwé, P. W. Schiller, B. L. Roth, S. Ballet, V. Katritch, R. C. Stevens and V. Cherezov (2015). “Structural basis for bifunctional peptide recognition at human δ-opioid receptor.” Nature structural & molecular biology 22(3): 265–268.

Ferreira, I. C., L. Barros, M. E. Soares, M. L. Bastos and J. A. Pereira (2007). “Antioxidant activity and phenolic contents of Olea europaea L. leaves sprayed with different copper formulations.” Food Chemistry 103(1): 188–195.

Gidudu, J., D. A. Sack, M. Pina, M. Hudson, K. Kohl, P. Bishop, A. Chatterjee, E. Chiappini, A. Compingbutra and C. Da Costa (2011). “Diarrhea: case definition and guidelines for collection, analysis, and presentation of immunization safety data.” Vaccine 29(5): 1053.

Guo-Hong, L., Z. Pei-Ji, S. Yue-Mao and Z. Ke-Qin (2004). “Antibacterial activities of neolignans isolated from the seed endotheliums of Trewia nudiflora.” Journal of Integrative Plant Biology 46(9): 1122–1127.

Gupta, K., A. Kumar, V. Tomer, V. Kumar and M. Saini (2019). “Potential of Colocasia leaves in human nutrition: Review on nutritional and phytochemical properties.” Journal of food biochemistry 43(7): e12878.

Hatano, T., T. Yasuhara, R. Yoshihara, I. Agata, T. Noro and T. Okuda (1990). “Effects of Interaction of Tannins with Co-existing Substances. VII.: Inhibitory Effects of Tannins and Related Polyphenols on Xanthine Oxidase.” Chemical and Pharmaceutical Bulletin 38(5): 1224–1229.

Horton, J. W. (2003). “Free radicals and lipid peroxidation mediated injury in burn trauma: the role of antioxidant therapy.” Toxicology 189(1-2): 75–88.

Hu, J., W.-Y. Gao, N.-S. Ling and C.-X. Liu (2009). “Antidiarrhoeal and intestinal modulatory activities of Wei-Chang-An-Wan extract.” Journal of ethnopharmacology 125(3): 450–455.

Ivancic, A., O. Roupsard, J. Q. Garcia, M. Melteras, T. Molisale, S. Tara and V. Lebot (2008). “Thermogenesis and flowering biology of Colocasia gigantea, Araceae.” Journal of plant research 121(1): 73–82.

Kähkönen, M. P., A. I. Hopia, H. J. Vuorela, J.-P. Rauha, K. Pihlaja, T. S. Kujala and M. Heinonen (1999). “Antioxidant activity of plant extracts containing phenolic compounds.” Journal of agricultural and food chemistry 47(10): 3954–3962.

Karaman, I., F. Şahin, M. Güllüce, H. Ögütçü, M. Şengül and A. Adıgüzel (2003). “Antimicrobial activity of aqueous and methanol extracts of Juniperus oxycedrus L.” Journal of ethnopharmacology 85(2-3): 231–235.

Khan, S., M. Nazir, N. Raiz, M. Saleem, G. Zengin, G. Fazal, H. Saleem, M. Mukhtar, M. I. Tousif, R. B. Tareen, H. H. Abdallah and F. M. Mahomoodally (2019). “Phytochemical profiling, in vitro biological properties and in silico studies on Caragana ambigua stocks (Fabaceae): A comprehensive approach.” Industrial Crops and Products 131: 117–124.

Konaté, K., K. Yomalan, O. Sytar and M. Brestic (2015). “Antidiarrheal and antimicrobial profiles extracts of the leaves from Trichilia emetica Vahl.(Meliaceae).” Asian Pacific Journal of Tropical Biomedicine 5(3): 242–248.

Kumaran, A. and R. J. Karunakaran (2007). “Activity-guided isolation and identification of free radical-scavenging components from an aqueous extract of Coleus aromaticus.” Food Chemistry 100(1): 356–361.

Kumaran, A. and R. J. Karunakaran (2007). “In vitro antioxidant activities of methanol extracts of five Phyllanthus species from India.” LWT-Food Science and Technology 40(2): 344–352.

Kuzuyama, T., J. P. Noel and S. B. Richard (2005). “Structural basis for the promiscuous biosynthetic prenylation of aromatic natural products.” Nature 435(7044): 983–987.

Lipinski, C. A., F. Lombardo, B. W. Dominy and P. J. Feeney (1997). “Experimental and computational approaches to estimate solubility and permeability in drug discovery and development settings.” Advanced Drug Delivery Reviews 23(1): 3–25.

Lipnick, R., J. Cotruvo, R. Hill, R. Bruce, K. Stitzel, A. P. Walker, I. Chu, M. Goddard, L. Segal and J. Springer (1995). “Comparison of the up-and-down, conventional LD50, and fixed-dose acute toxicity procedures.” Food and chemical toxicology 33(3): 223–231.

Liu, B., Y. Liu, W. Cao, S. Zhang, Z. Liu, Y. Ni and F. Li (2014). “Ethnobotany of Medicinal Aroids in Xishuangbanna, Yunnan Province, China.” Aroideana: 69.

Lutterodt, G. D. (1992). “Inhibition of Microlax-induced experimental diarrhoea with narcotic-like extracts of Psidium guajava leaf in rats.” Journal of Ethnopharmacology 37(2): 151–157.

Mahboubi, M. and G. Haghi (2008). “Antimicrobial activity and chemical composition of Mentha pulegium L. essential oil.” Journal of ethnopharmacology 119(2): 325–327.

Maridass, M. and A. J. De Britto (2008). “Origins of plant derived medicines.” Ethnobotanical Leaflets 2008(1): 44.

Mayo, S., J. Bogner and P. Boyce (1997). “The Genera of Araceae Royal Botanical Gardens.” London, uK.

Mbagwu, H. and O. Adeyemi (2008). “Anti-diarrhoeal activity of the aqueous extract of Mezoneuron benthamianum Baill (Caesalpiniaceae).” Journal of Ethnopharmacology 116(1): 16–20.

Meite, S., J. N‘guessan, C. Bahi, H. Yapi, A. Djaman and F. G. Guina (2009). “Antidiarrheal activity of the ethyl acetate extract of Morinda morindoides in rats.” Tropical Journal of Pharmaceutical Research 8(3).

Mendonça-Filho, R. R. (2006). “Bioactive phytocompounds: new approaches in the phytosciences.” Modern phytomedicine: Turning medicinal plants into drugs.

Mishra, A., A. Seth and S. K. Maurya (2016). “Therapeutic significance and pharmacological activities of antidiarrheal medicinal plants mention in Ayurveda: A review.” Journal of intercultural ethnopharmacology 5(3): 290.

Mukul, S. A., M. B. Uddin and M. R. Tito (2007). “Medicinal plant diversity and local healthcare among the people living in and around a conservation area of Northern Bangladesh.” International Journal of Forest Usufructs Management 8(2): 50–63.

Muramatsu, H., K. Kogawa, M. Tanaka, K. Okumura, Y. Nishihori, K. Koike, T. Kuga and Y. Niitsu (1995). “Superoxide dismutase in SAS human tongue carcinoma cell line is a factor defining invasiveness and cell motility.” Cancer Research 55(24): 6210–6214.

Novakova, Z., J. Cerny, C. J. Choy, J. R. Nedrow, J. K. Choi, J. Lubkowski, C. E. Berkman and C. Barinka (2016). “Design of composite inhibitors targeting glutamate carboxypeptidase II: the importance of effector functionalities.” The FEBS journal 283(1): 130–143.

Oladeji, O. (2016). “The characteristics and roles of medicinal plants: Some important medicinal plants in Nigeria.” Indian Journal of Natural Products 12(3): 102.

Page, A.-L., S. Hustache, F. J. Luquero, A. Djibo, M. L. Manzo and R. F. Grais (2011). “Health care seeking behavior for diarrhea in children under 5 in rural Niger: results of a cross-sectional survey.” BMC public health 11(1): 389.

Petrovska, B. B. (2012). “Historical review of medicinal plants’ usage.” Pharmacognosy reviews 6(11): 1.

Prlic, A., S. Bliven, P. W. Rose, W. F. Bluhm, C. Bizon, A. Godzik and P. E. Bourne (2010). “Pre-calculated protein structure alignments at the RCSB PDB website.” Bioinformatics 26(23): 2983–2985.

Qin, Y., J.-b. Wang, W.-j. Kong, Y.-l. Zhao, H.-y. Yang, C.-m. Dai, F. Fang, L. Zhang, B.-c. Li and C. Jin (2011). “The diarrhoeogenic and antidiarrhoeal bidirectional effects of rhubarb and its potential mechanism.” Journal of ethnopharmacology 133(3): 1096–1102.

Qiu, X., C. A. Janson, W. W. Smith, M. Head, J. Lonsdale and A. K. Konstantinidis (2001). “Refined structures of beta-ketoacyl-acyl carrier protein synthase III.” J Mol Biol 307(1): 341–356.

Saeed, N., M. R. Khan and M. Shabbir (2012). “Antioxidant activity, total phenolic and total flavonoid contents of whole plant extracts Torilis leptophylla L.” BMC complementary and alternative medicine 12(1): 221.

Sher, A. (2009). “Antimicrobial activity of natural products from medicinal plants.” Gomal Journal of Medical Sciences 7(1).

Shiferie, F. and W. Shibeshi (2013). “In vivo antidiarrheal and ex-vivo spasmolytic activities of the aqueous extract of the roots of Echinops kebericho Mesfin (Asteraceae) in rodents and isolated Guinea-pig ileum.” Int J Pharm Pharmacol 2: 110–116.

Sini, J., I. Umar, K. Anigo, I. Stantcheva, E. Bage and R. Mohammed (2008). “Antidiarrhoeal activity of aqueous extract of Combretum sericeum roots in rats.” African journal of Biotechnology 7(17).

Steinberg, D., S. Parthasarathy, T. E. Carew, J. C. Khoo and J. L. Witztum (1989). “Beyond cholesterol.” New England Journal of Medicine 320(14): 915–924.

Tunaru, S., T. F. Althoff, R. M. Nüsing, M. Diener and S. Offermanns (2012). “Castor oil induces laxation and uterus contraction via ricinoleic acid activating prostaglandin EP3 receptors.” Proceedings of the National Academy of Sciences 109(23): 9179–9184.

Wagner, W., D. Herbst and S. Sohmer (1990). “Manual of the flowering plants of Hawai =i. University of Hawai =i Press.” Bishop Museum Special Publication 83.

Wansi, S. L., E. P. Nguelefack-Mbuyo, M. L. Nchouwet, D. Miaffo, P. Nyadjeu, J. P. Wabo, M. Mbiantcha, P. A. NKeng-Efouet, T. B. Nguelefack and A. Kamanyi (2014). “Antidiarrheal activity of aqueous extract of the stem bark of Sapium Ellipticum (Euphorbiaceae).” Tropical Journal of Pharmaceutical Research 13(6): 929–935.

Wu, H., D. Wacker, M. Mileni, V. Katritch, G. W. Han, E. Vardy, W. Liu, A. A. Thompson, X.-P. Huang, F. I. Carroll, S. W. Mascarella, R. B. Westkaemper, P. D. Mosier, B. L. Roth, V. Cherezov and R. C. Stevens (2012). “Structure of the human κ-opioid receptor in complex with JDTic.” Nature 485(7398): 327–332.

Yadav, J., S. Kumar and P. Siwach (2006). “Folk medicine used in gynecological and other related problems by rural population of Haryana.”

Yin, J.-T. (2006). Colocasia tibetensis (Araceae, Colocasieae), a new species from southeast Tibet, China. Annales Botanici Fennici, JSTOR.

Zhang, J., S. Wang, Y. Li, P. Xu, F. Chen, Y. Tan and J. Duan (2013). “Anti-diarrheal constituents of Alpinia oxyphylla.” Fitoterapia 89: 149–156.

Zheng, W. and S. Y. Wang (2001). “Antioxidant activity and phenolic compounds in selected herbs.” Journal of Agricultural and Food chemistry 49(11): 5165–5170.

